# The role of the fornix in human navigational learning

**DOI:** 10.1101/391888

**Authors:** Carl J. Hodgetts, Martina Stefani, Angharad N. Williams, Branden S. Kolarik, Andrew P. Yonelinas, Arne D. Ekstrom, Andrew D. Lawrence, Jiaxiang Zhang, Kim S. Graham

## Abstract

Studies in rodents have demonstrated that transecting the white matter pathway linking the hippocampus and anterior thalamic nuclei - the fornix - impairs flexible navigational learning in the Morris Water Maze (MWM), as well as similar spatial learning tasks. While diffusion MRI studies in humans have linked fornix microstructure to scene discrimination and memory, its role in human navigation is currently unknown. We used high-angular resolution diffusion MRI to ask whether inter-individual differences in fornix microstructure would be associated with spatial learning in a virtual MWM task. To increase sensitivity to individual learning across trials, we adopted a novel curve fitting approach to estimate a single index of learning rate. We found a significant correlation between learning rate and the microstructure (mean diffusivity) of the fornix, but not that of a control tract linking occipital and anterior temporal cortices (the inferior longitudinal fasciculus, ILF). Further, this correlation remained significant when controlling for hippocampal volume. These findings extend previous animal studies by demonstrating the functional relevance of the fornix for human navigational learning, and highlight the importance of a distributed neuroanatomical network, underpinned by key white matter pathways, such as the fornix, in complex spatial behaviour.

## Introduction

The ability to navigate, and learn the location of rewards and goals in the environment, is a fundamental and highly adaptive cognitive function across species (Landau and Lakusta, 2009; Wolbers and Hegarty, 2010; Murray et al., 2016). Lesion studies in animals suggest that this ability depends, in part, on several key brain regions, including the hippocampus, mammillary bodies, and the anterior thalamic nuclei (Sutherland and Rodriguez, 1989; Warburton and Aggleton, 1998; Jankowski et al., 2013), which in turn connect with a broader network including entorhinal, parahippocampal, retrosplenial, and posterior parietal cortex, all thought to be important for navigation (Ekstrom et al., 2017). In particular, the hippocampus, mammillary bodies, and anterior thalamic nuclei are connected anatomically by an arch-shaped white matter pathway called the fornix (Saunders and Aggleton, 2007). Given the role of these interconnected structures in spatial learning and navigation (Jankowski et al., 2013), the ability for these distributed regions to communicate via the fornix may also be critical for successful spatial learning and navigation.

Indeed, transecting the fornix in rodents and monkeys impairs learning for objects-in-place, but not the objects themselves (Gaffan, 1992, 1994; Simpson et al., 1998). These findings also extend to performance on spatial navigation tasks, most notably the Morris Water Maze (MWM). The MWM is one of the most widely used laboratory tasks in studies of navigational behaviour across non-human species and has been recognized as an excellent candidate for a universal test of spatial navigation ability (Morris, 1984; Possin et al., 2016). In this task, animals are placed in a circular pool and required to swim to a hidden platform beneath the surface using allocentric cues outside the pool. Several studies have shown that fornix-transected rodents are impaired on the MWM, particularly when required to navigate flexibly from multiple positions within the maze (Eichenbaum et al., 1990; Packard and McGaugh, 1992; Warburton et al., 1998; Warburton and Aggleton, 1998; De Bruin et al., 2001; Cain et al., 2006). Fornix transection also impairs allocentric place learning in other maze tasks (O’Keefe et al., 1975; Olton et al., 1978; Packard et al., 1989; Dumont et al., 2015).

Critically, while these animal studies highlight a key role for the fornix in spatial learning - across both visuo-spatial discrimination and navigation tasks - the role of this white matter pathway in human wayfinding is currently unknown. Studies using diffusion magnetic resonance imaging (dMRI), which allows white matter microstructure to be quantified *in vivo*, have reported associations in healthy human subjects between fornix microstructure and inter-individual differences in scene and spatial context processing across both memory (Rudebeck et al., 2009; Hodgetts et al., 2017) and perceptual tasks (Postans et al., 2014; Hodgetts et al., 2015). Given differences in the visuospatial representations underpinning navigation across rodents and humans (Ekstrom, 2015), it begs the question whether this same extended functional system, structurally linked by the fornix, is similarly important for navigational learning in humans.

To test this, we acquired dMRI data in healthy human subjects who performed a human analogue of the MWM (Figure 1). In this task, individuals were required to learn, over trials, the location of a hidden sensor within a virtual art gallery. Similar to the rodent paradigm, subjects were required to navigate from multiple starting positions, thus placing greater demand on flexible allocentric processing (Figure 1). To create a single index of navigational learning rate, we used a curve fitting approach to model the time taken to reach the sensor across trials (for similar approaches, see Stepanov and Abramson, 2008; Pereira and Burwell, 2015; Kahn et al., 2017). We predicted, based on previous work (Packard and McGaugh, 1992; Warburton and Aggleton, 1998; Cain et al., 2006; Hodgetts et al., 2015), that microstructure of the fornix, but not a control tract connecting occipital and anterior temporal cortices (the “inferior longitudinal fasciculus”, ILF) (Latini, 2015), would be significantly related to spatial learning rate in a virtual MWM task.

**Figure 1.**
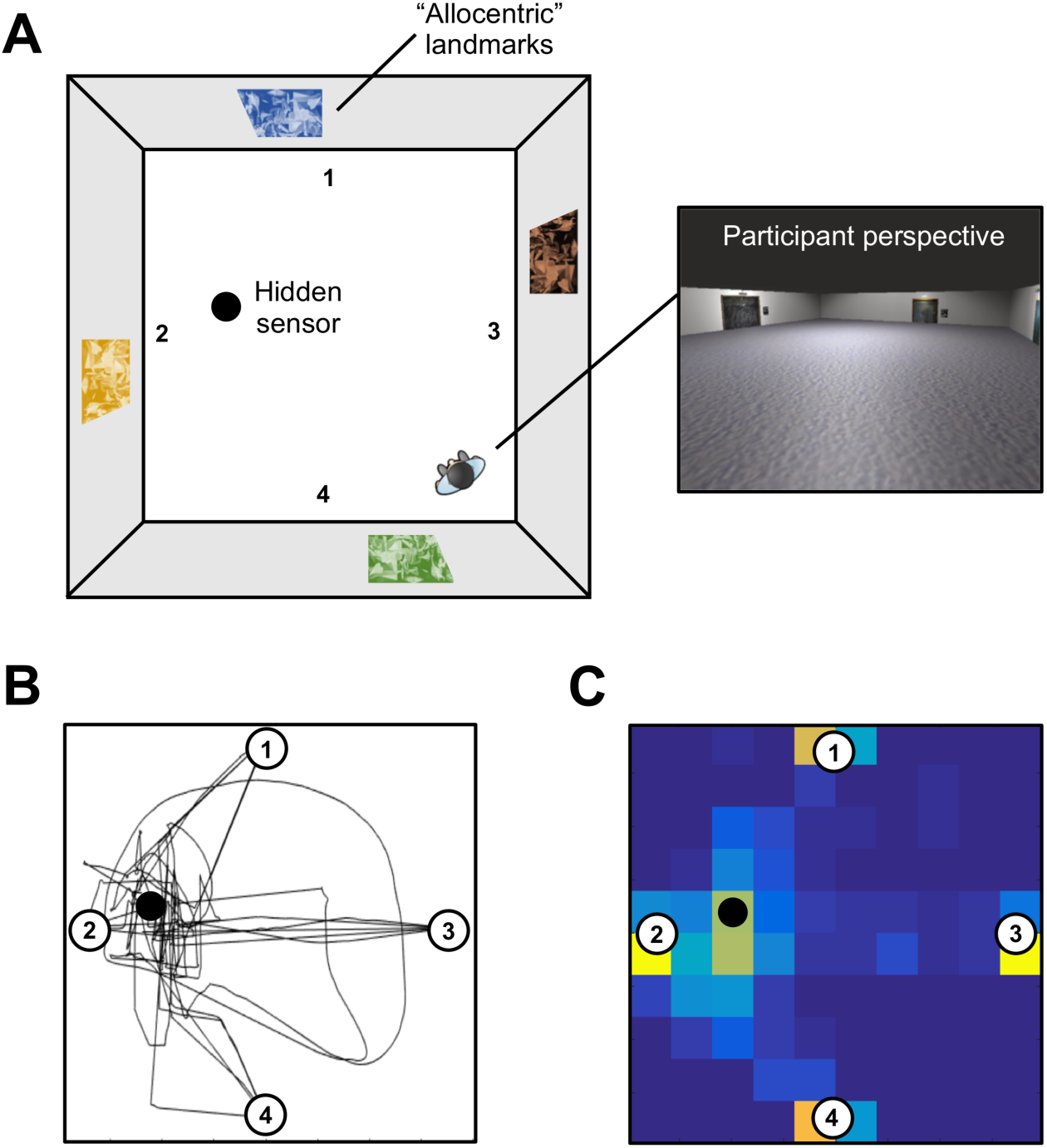
The virtual reality Morris Water Maze. (A) Birds-eye schematic of the virtual art gallery that the participants explore during the task. The artwork on the outer walls of the gallery are the “landmarks” in the virtual arena. An example first person perspective from within the maze is shown. (B) Movement trajectories and (C) location heatmap across all 20 trials for an example participant.

## Methods

### Participants

Thirty-three healthy volunteers (15 males, 18 females; mean age = 24 years; SD = 3.5 years) were scanned at the Cardiff University Brain Research Imaging Centre (CUBRIC). These same participants completed a virtual Morris Water Maze task in a separate behavioural session. All subjects were fluent English speakers with normal or corrected-to-normal vision. Participation in both sessions was undertaken with the understanding and written consent of each subject. The research was completed in accordance with, and approved by, the Cardiff University School of Psychology Research Ethics Committee.

### Virtual Morris Water Maze Task

We used the virtual MWM task developed by Kolarik et al. (2016). This task was created using Unity 3D (Unity Technologies, San Francisco) and required participants to explore, from a first-person perspective, a virtual art gallery using the arrow keys on the computer keyboard (Figure 1A). The room was 8 x 8 virtual m^2^ in size, and contained four distinct paintings, one on each wall of the environment. On a given trial, the participants’ task was to locate a hidden sensor on the floor as quickly as possible. This sensor occupied 0.25% of the total floor space (i.e., an 0.8 x 0.8 m^2^ square). When the participant walked over the hidden platform it became visible and the caption ‘You found the hidden sensor’ was displayed in the centre of the screen. At this point, the exploration time was recorded automatically and a 10 second countdown appeared in the centre of the display during which the participants could freely navigate the room. After this countdown, an inter-trial window appeared and the participants could click on a button to start the next learning trial. The maximum duration of each learning trial was 60 seconds. If the participant did not find the target location within this period, the sensor became visible. The task involved 20 learning trials, which comprised five blocks of four trials. Within each block, participants started from each of the four starting positions (arbitrary North, South, East, West). The movement trajectories and location heatmap for an example participant is shown in Figure 1B-C.

### MRI acquisition

Whole brain dMRI data were acquired at the Cardiff University Brain Research Imaging Centre (CUBRIC) using a 3T GE HDx Signa scanner with an eight-channel head coil. Single-shell high-angular resolution dMRI (HARDI) (Tuch et al., 2002) data were collected with a single-shot spin-echo echo-planar imaging pulse sequence with the following parameters: 30 directions; TE= 87 ms; 60 continuous slices acquired along an oblique-axial plane with 2.4 mm thickness and no gap. The scans were cardiac-gated using a peripheral pulse oximeter placed on the participants’ fingertips. A T1-weighted 3D FSPGR sequence was also acquired with the following parameters: TR= 7.8 ms; TE= 3 ms, TI= 450 ms, flip angle= 20°; FOV= 256 mm*192 mm*172 mm; 1 mm isotropic resolution.

### Diffusion MRI preprocessing

Diffusion MRI data were corrected for subject head motion and eddy currents using ExploreDTI (Version 4.8.3; Leemans and Jones, 2009). The bi-tensor ‘Free Water Elimination’ (FWE) procedure was applied *post hoc* to correct for voxel-wise partial volume artifacts arising from free water contamination (Pasternak et al., 2009). Free water contamination (from cerebrospinal fluid) is a particular issue for white matter pathways located near the ventricles (such as the fornix), and has been shown to significantly affect tract delineation (Concha et al., 2005). Following FWE, corrected diffusion-tensor indices FA and MD were computed. FA reflects the extent to which diffusion within biological tissue is anisotropic, or constrained along a single axis, and can range from 0 (fully isotropic) to 1 (fully anisotropic). MD (10^-3^mm^2^s^-1^) reflects a combined average of axial diffusion (diffusion along the principal axis) and radial diffusion (diffusion along the orthogonal direction).

### Tractography

Deterministic whole brain white matter tractography was performed using the ExploreDTI graphical toolbox. Tractography was based on constrained spherical deconvolution (CSD) (Jeurissen et al., 2011), which can extract multiple peaks in the fiber orientation density function (fODF) at each voxel. This approach permits the representation of crossing/kissing fibers in individual voxels. Each streamline was reconstructed using an fODF amplitude threshold of 0.1 and a step size of 1mm, and followed the peak in the fODF that subtended the smallest step-wise change in orientation. An angle threshold of 30° was used and any streamlines exceeding this threshold were terminated.

Three-dimensional reconstructions of each tract were obtained from individual subjects by using a waypoint region of interest (ROI) approach, based on an anatomical prescription. Here, “AND” and “NOT” gates were applied, and combined, to extract tracts from each subject’s whole brain tractography data. These ROIs were drawn manually on the direction-encoded FA maps in native space by one experimenter (MS) and quality assessed by other experimenters (CJH, ANW).

### Fornix

A multiple region-of-interest (ROI) approach was adopted to reconstruct the fornix (Metzler-Baddeley et al., 2011). This approach involved placing a seed point ROI on the coronal plane at the point where the anterior pillars enter the fornix body. Using a mid-sagittal plane as a guide, a single AND ROI was positioned on the axial plane, encompassing both crus fornici at the lower part of the splenium of the corpus callosum. Three NOT ROIs were then placed: (1) anterior to the fornix pillars; (2) posterior to the crus fornici; and (3) on the axial plane, intersecting the corpus callosum. Once these ROIs were placed, and the tracts reconstructed, anatomically implausible fibers were removed using additional NOT ROIs (see Hodgetts et al., 2017).

### Inferior longitudinal fasciculus (ILF)

Fiber-tracking of the ILF (control tract) was performed using a two-ROI approach in each hemisphere (Wakana et al., 2007). First, the posterior edge of the cingulum bundle was identified on the sagittal plane. Reverting to a coronal plane at this position, a SEED ROI was placed that encompassed the whole hemisphere. To isolate streamlines extending towards the anterior temporal lobe (ATL), a second ROI was drawn at the most posterior coronal slice in which the temporal lobe was not connected to the frontal lobe. Here, an additional AND ROI was drawn around the entire temporal lobe. Similar to the fornix protocol above, any anatomically implausible streamlines were removed using additional NOT ROIs. This approach was carried out in both hemispheres; diffusion properties of the left and right ILF (for both FA and MD) were averaged across hemispheres to provide a bilateral measure of ILF FA and MD in each participant.

### Grey matter volumetry

Bilateral hippocampal volume was derived using FMRIB’s Integrated Registration & Segmentation Tool (FIRST; Patenaude et al., 2012). As temporal lobe substructures have been shown to correlate with intracranial volume (Moran et al., 2001), individual-level hippocampal volumes were divided by total intracranial volume (eTIV) to create proportional scores (Westman et al., 2013).

### Statistical analysis of maze learning

To increase sensitivity to individual-level performance across learning trials, and to derive a single index of learning rate, we analysed the relationship between spatial learning and fornix tissue microstructure using a curve fitting approach (see e.g., Pereira and Burwell, 2015; Kahn et al., 2017). Performance on each learning trial was defined by the time (in seconds) to reach the hidden sensor. As can be seen in Figure 2A, there was high inter-individual variability in spatial learning, with subjects varying in both learning speed and the shape of their learning pattern. Here, individual learning data was fit using a power function: Time to sensor = a * xb, where b specifies the slope of the fitted power model.

**Figure 2.**
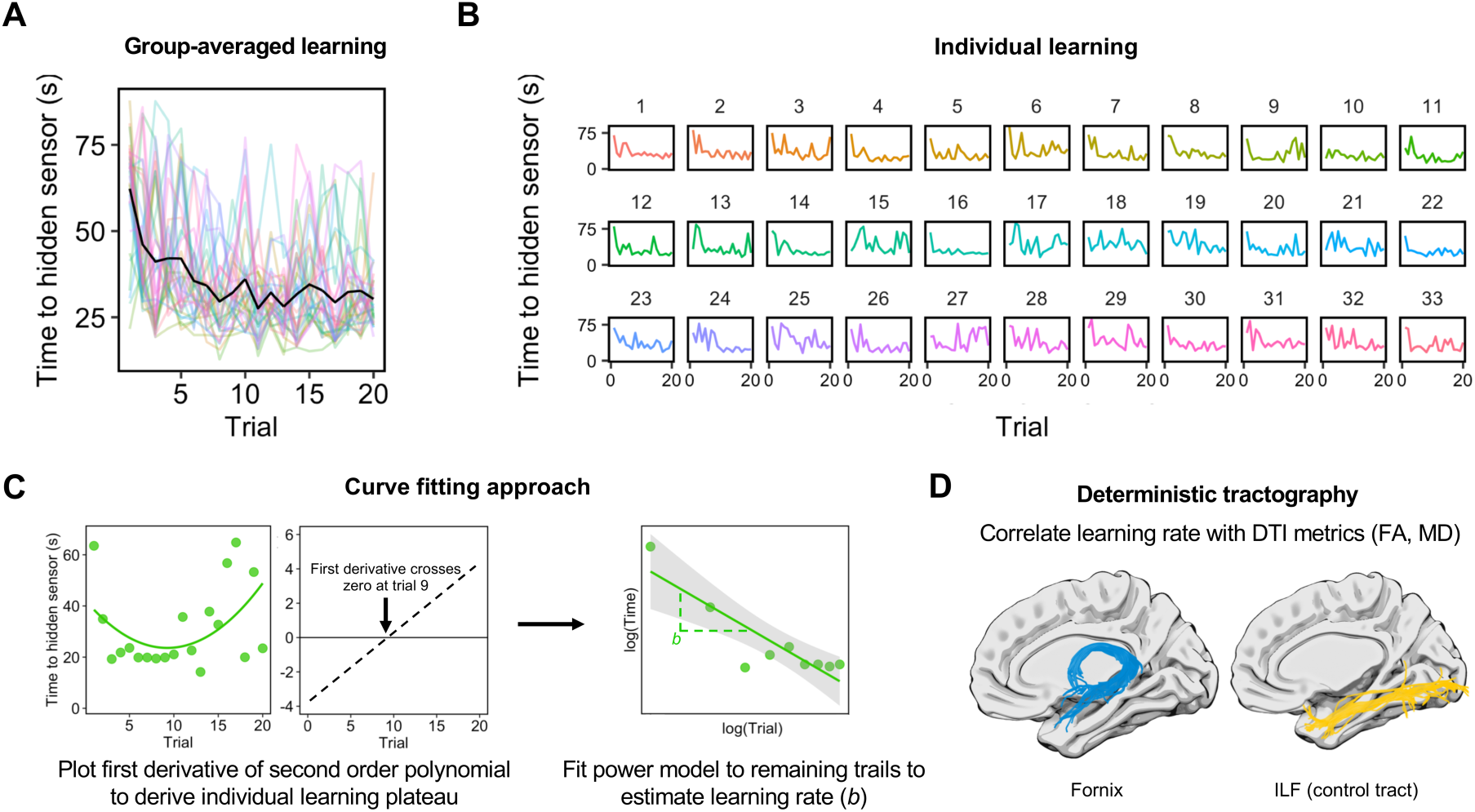
Modelling navigational learning in individual participants. MWM task learning at the (A) group-level and (B) individual-level. Y-axes represent the time to reach the hidden sensor in seconds. The number of trials (total = 20) is shown on the x-axis. (C) Method for determining the number of learning trials to-be-modelled. Some participants appeared to learn rapidly and plateau before displaying variable performance in later trials. For instance, a power model fits the example participant’s latency data poorly when all trials are considered. In order to capture initial learning, therefore, we fitted the latency data (across all trials) with a second-order polynomial in each subject. The point at which the first derivative of this polynomial crossed zero was used to define the number of trials to-be-modelled. The trials up to this point were then fit with a power function and the b parameter derived to index learning rate. Power fits are shown by linearly fitting the log-transformed data. (D) Learning rate measures were correlated with diffusion metrics (FA, MD) from the fornix (blue) and the ILF (yellow). Tract reconstructions are shown against an inflated brain for visualisation purposes.

One aspect of this data is that some subjects learned quickly (and plateaued) before displaying variable, or slow, performance in the later trials (e.g., subjects 9, 13, and 20; Figure 2B). This presents a challenge for a curve fitting approach across all trials (and potentially produces counterintuitive results), as some of the fastest learners will show the poorest model fits. For instance, both subjects 9 and 16 display an initial steep learning curve and an early plateau (Figure 2B), but a power model fit to *all trials* provides a poor fit of the subject who does not sustain performance until the end of the task. In order to account for this complexity in learning patterns, we adopted a data-driven approach to determine a cut-off in individual subjects. Specifically, a second-order polynomial model was fit to all trials in each subject using the curve fitting toolbox in Matlab (Mathworks, Inc.). The cut-off was defined as the trough of this curve, which is where the first derivative of the second-degree polynomial crosses zero (Figure 2C). Trials up to and including this cut-off were then modelled using a power function (mean trials included = 14.3; range = 7 – 20).

Using this approach, we derived a single measure of learning rate, denoted by the b parameter (or slope) of the fitted power model (b; mean = -0.32, SD = 0.08, range = -0.49 to -0.19). The b parameter reflects slope curvilinearity in each subject, where lower, negative values reflect more convex downward curves and thus faster learning rates. As such, we predict a positive association between fornix MD and learning rate, and negative associations between fornix FA and learning rate.

Directional Pearson correlations were conducted between the learning rate and free water corrected MD and FA values for the fornix and ILF (Figure 2D). The resulting coefficients were compared statistically using directional Steiger Z-tests (Steiger, 1980) within the ‘cocor’ package in R (Diedenhofen and Musch, 2015). Pearson correlations were Bonferroni-corrected by dividing α = 0.05 by the number of statistical comparisons for each DTI metric (i.e., 0.05/2 = 0.025) (Lakens, 2016). Prior to correlational analyses, outliers for each tract and metric were identified and removed using the Tukey method in R. This excluded an extreme value for fornix MD, fornix FA, and ILF FA. To exclude poor performers who were not engaging with the task, we used a resampling approach where individual-level data was shuffled over 500 permutations and confidence intervals (CIs) derived. Participants with a model R^2^ that fell outside the CI of their individually-defined random distribution were excluded (Subjects 10, 15, 17, 18 and 21).

We also conducted Bayesian correlation analyses using JASP (https://jasp-stats.org). From this, we report default Bayes factors and 95% Bayesian credibility intervals (BCI). The Bayes factor, expressed as BF_10_ grades the intensity of the evidence that the data provide for the alternative hypothesis (H1) versus the null (H0) on a continuous scale. A BF_10_ of 1 indicates that the observed finding is equally likely under the null and the alternative hypothesis. A BF_10_ much greater than 1 allows us to conclude that there is substantial evidence for the alternative over the null. Conversely BF_10_ values substantially less than 1 provide strong evidence in favour of the null over the alternative hypothesis (Wetzels and Wagenmakers, 2012).

Complementary Spearman’s rho tests were also conducted for our key correlations. The strength of Spearman’s correlations were compared directly using a robust bootstrapping approach (Wilcox, 2016), as implemented using ‘comp2dcorr’ in Matlab (https://github.com/GRousselet/blog/tree/master/comp2dcorr).

## Results

### Correlating navigational learning with tract microstructure

There was a significant positive correlation between the derived learning rate and fornix MD, as shown in Figure 3. This suggests that those subjects with lower fornix MD had faster learning rates (r = 0.44, p = 0.01, 95% BCI [0.09, 0.68], B_+0_ = 5.5; Figure 3). There was no significant relationship between individual learning rate and MD in a control tract - the inferior longitudinal fasciculus (ILF; r = -0.06; p = 0.62, 95% BCI [0.37, 0.01], B_0+_ = 5.38). A directional Steiger Z-test (Steiger, 1980) revealed that the correlation between derived learning rate and fornix MD was significantly greater than with ILF MD (z = 2.26, p = 0.01).

**Figure 3.**
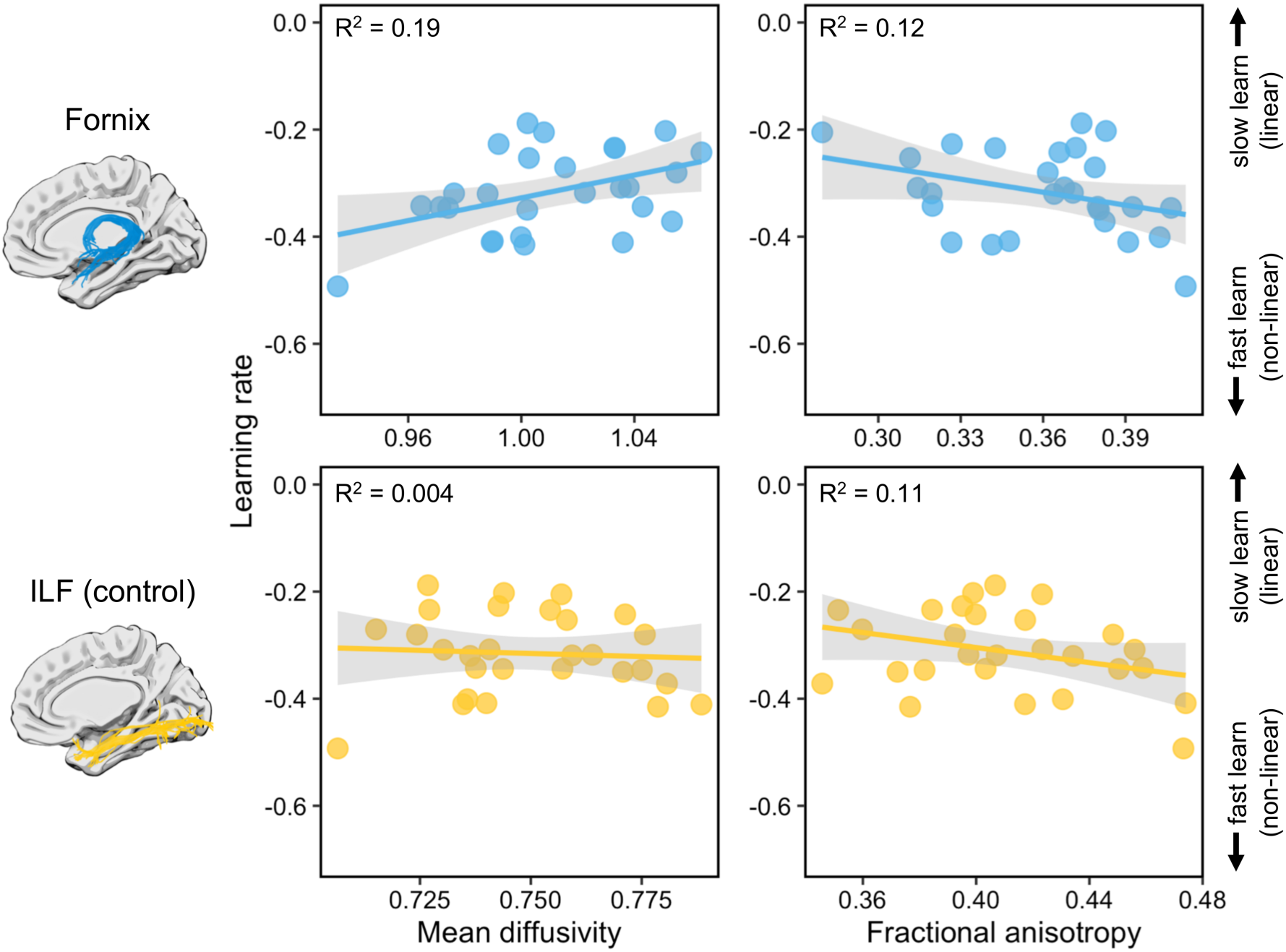
The correlation between tract microstructure and learning rate (b parameter) for the fornix (top row) and the inferior longitudinal fasciculus (ILF).

A moderate trend was observed between fornix FA and learning rate but this did not reach our experiment-wise significance level (r = -0.34, p = 0.04, 95% BCI [-0.62, - 0.04], B_-0_ = 1.99; Figure 3). There was no significant correlation between ILF FA and learning rate (r = -0.17; p = 0.2, 95% BCI [-0.51, -0.01], B_-0_ = 1.68). These two correlations did not differ significantly (z = 0.22, p = 0.21).

### Controlling for hippocampal volume

To examine whether hippocampal volume contributes to the microstructural-behavioural correlations reported above, partial correlations (both frequentist and Bayesian) were conducted. The significant positive correlation between the learning rate parameter and fornix MD remained when controlling for bilateral hippocampal volume (r = 0.4, p = 0.02, BF_+0_ = 3.59), as seen in prior studies (Hodgetts et al., 2017). For fornix FA, a slightly stronger negative trend was observed (r = -0.35, p = 0.04, BF_-0_ = 0.09) when hippocampal volume was controlled for, though this did not reach our experiment-wise significance level (i.e., p = 0.025). When examining hippocampal volume, independent of fornix microstructural measures, there was no significant association found between hippocampal volume and learning rate (r = 0.03, p = 0.94, 95% BCI [-0.25, -0.002], B_-0_ = 10.2).

### Non-parametric correlations between tract microstructure and learning

Finally, we also conducted complementary directional Spearman’s rho tests for our key correlations, with such tests robust to univariate outliers (Croux and Dehon, 2010). As above, Spearman’s correlations were Bonferroni-corrected by dividing α = 0.05 by the number of statistical comparisons for each DTI metric (i.e., 0.05/2 = 0.025). A significant positive association was observed between learning rate and fornix MD (ρ = 0.4, p = 0.02). No significant association was found with ILF MD (ρ = -0.18, p = 0.82). A strong trend was found between the b parameter and fornix FA (ρ = -0.32, p = 0.05) but not ILF FA (ρ = -0.21, p = 0.14).

A direct comparison between these correlations revealed a significant difference between fornix MD and ILD MD and their association with navigation learning rate, as indicated by the bootstrap distribution not overlapping with zero (95% CI = 0.2 – 0.88, p = 0). There was no significant difference between the FA correlations (95% CI = -0.7191 - 0.2962, p = 0.4).

## General discussion

Using a virtual-reality analogue of a classic navigational paradigm, the Morris Water Maze (Morris, 1984), we asked whether inter-individual variation in the microstructure of the fornix (linking hippocampus with medial diencephalon and prefrontal cortex) is related to individual differences in navigational learning. To increase sensitivity to individual learning across trials we adopted a curve fitting approach (Kahn et al., 2017), which generated a single index of learning rate (‘b’) in each individual. We found that fornix microstructure (particularly MD) was significantly associated with navigational learning rate in a virtual MWM task, as defined by the slope of the fitted power model, and this association remained when controlling for bilateral hippocampal volume. Furthermore, this effect was significantly stronger than that seen for the ILF, a control tract linking occipital and anterior temporal cortices, which has previously been implicated in semantic learning (Qi et al., 2015; Ripollés et al., 2017).

These results build upon previous animal studies that highlight a potential key role for the fornix in mediating place learning and navigational behaviour. Critically, we provide novel evidence, using a MWM task analogous to that used in animals (Kolarik et al., 2016; Possin et al., 2016), that the fornix supports navigational learning in humans. In rodents, fornix transection has been shown to impair MWM learning, as characterised by more gradual learning slopes and slower latencies in finding the hidden platform (Eichenbaum et al., 1990; Packard and McGaugh, 1992; Warburton and Aggleton, 1998; Cain et al., 2006). By applying a curve fitting approach, we were able to characterise the steepness of learning slopes at the individual participant level, and relate this directly with fornix microstructure. Strikingly consistent with the animal studies described above, reduced structural connectivity in the fornix (indexed by higher MD) was related to more gradual learning rates. Further, by identifying individual learning plateaus in a data-driven way, this approach also accounts for potential fatigue, mind-wandering or other factors that may affect performance later in the learning session.

Similar to lesioning hippocampus and anterior thalamic nuclei, learning deficits following fornix transection in rodents are also more severe when the animal is required to navigate from multiple start positions (Eichenbaum et al., 1990). Such findings suggest, therefore, that this broader neuroanatomical system, structurally underpinned by the fornix (Aggleton et al., 2010), supports spatial learning in a flexible manner (i.e., from novel start positions, or from different perspectives), rather than response-based learning, that appears to recruit regions outside this extended hippocampal system, specifically the caudate nucleus (Packard and McGaugh, 1992; Devan et al., 1996; Chersi and Burgess, 2015). Consistent with this, we observed an association between navigational learning and fornix properties in a task which required participants to navigate to the goal from multiple starting positions.

Overall, this study provides support for the idea that an individual’s spatial navigation ability (Wolbers and Hegarty, 2010) is underpinned, at least in part, by the integrated functioning of a distributed neuroanatomical network, comprising not only individual regions (such as the hippocampus and anterior thalamic nuclei), but also the white matter connections linking these brain areas (Jankowski et al., 2013; Murray et al., 2016). While MWM performance is considered to depend, at least partly, on the ability to form and utilise detailed allocentric mental representations, or “cognitive maps” (Tolman, 1948; O’Keefe and Nadel, 1976), human and animal studies suggest that the role of the fornix in spatial processing may be linked to mechanisms beyond spatial mapping *per se*.

For instance, while fornix transection impairs, or at least slows, navigational learning in the MWM (Warburton and Aggleton, 1998), as discussed above, these impairments are not as severe as that seen following lesions to the anterior thalamic nuclei or the hippocampus proper (Eichenbaum et al., 1990; Warburton and Aggleton, 1998; Cain et al., 2006). This is not to suggest that fornix connectivity is not important for place representations (Miller and Best, 1980; Shapiro et al., 1989), but rather that the fornix may support processes which help build and support detailed cognitive maps (e.g., scene-based processing, path integration) in conjunction with other brain areas involved in a broader navigation network (Whishaw and Maaswinkel, 1998; Gaffan et al., 2001). For instance, evidence from non-human primates suggests a potential key role in forming conjunctive scene representations (Gaffan, 1991; Hodgetts et al., 2015; Murray et al., 2017). The ability to learn and remember object-in-scene associations, as well as naturalistic scenes, is impaired significantly following fornicectomy (Gaffan, 1992; Gaffan et al., 2001; Buckley et al., 2008).

Convergent with scene learning deficits reported in monkeys, diffusion MRI studies in humans have reported associations between fornix microstructure and scene recollection (Rudebeck et al., 2009), complex scene discrimination (Postans et al., 2014; Hodgetts et al., 2015) and the ability to retrieve spatiotemporal detail in real-world memories (Hodgetts et al., 2017). Rather than suggesting a selective role in allocentric spatial navigation *per se*, these studies support the view that the connections established by the fornix may be critical for integrating scenes into coherent spatial representations, which then may contribute to the generation of detailed map-like representations useful for navigation (Ryan et al., 2010; Fidalgo and Martin, 2016). An alternative account (Relational Memory Theory), by contrast, posits that while the extended hippocampal system is essential to spatial navigation via a cognitive map, its role derives from the relational organization and flexibility of cognitive maps and not from a selective role in the spatial domain (Eichenbaum, 2017; see also Ekstrom and Ranganath, 2017). The initial formation of such flexible spatial relations has been argued to critically rely on cholinergic system modulation of the hippocampus (Ikonen et al., 2002), which is dependent on the fornix (Alonso et al., 1996), consistent with our findings.

Note, it is possible that some individual differences in navigational performance may actually reflect differences in types of spatial strategies employed. For instance, while some individuals may use a strategy akin to cognitive mapping, i.e., based on allocentric vectors from the “landmarks” to the hidden sensor, some individuals may use a strategy based on matching and integrating disparate viewpoints from the sensor location; a strategy more akin to building a model of the broader scene and layout (Wolbers and Wiener, 2014). While participants were not asked about their use of spatial strategies in the current study, this would be an interesting avenue for disentangling scene-based and cognitive mapping approaches in future studies.

While our findings support the notion that an extended hippocampal-based system, mediated by the fornix, may be important for navigational learning in humans, it was notable that the fornix association was present when controlling for HC volume. Further, there was no independent association between place learning and HC volume in this task. Though some studies have found associations between hippocampal grey matter volume and navigational ability in humans (Maguire et al., 1997; Bohbot et al., 1998; Schinazi et al., 2013; Chrastil et al., 2017), others have shown that fornix microstructure (but not hippocampal volume) predicts individual differences in remembering spatiotemporal aspects of autobiographical memories (e.g., Hodgetts et al., 2017). In addition, studies of individuals with profound orientation deficits (termed development topographical disorientation, or DTD) similarly show impairments in connectivity patterns to the hippocampus (in this case, between hippocampus and prefrontal cortex). Interestingly, like in our study, hippocampal grey matter does not appear to explain these differences (Iaria et al., 2009; Iaria and Barton, 2010). This highlights that variation in broader neuroanatomical systems, rather than regional volumetric variation, may be particularly sensitive to individual differences in navigational learning.

Similar to our previous work, we observed stronger effects for fornix MD versus FA (Postans et al., 2014; Hodgetts et al., 2015). The biological interpretation of this difference is not straightforward, as variation in either measure could arise from multiple aspect(s) of the underlying white matter, including axon density, axon diameter, myelination, and the manner in which fibres are arranged in a voxel (Beaulieu, 2002). A recent study reported strong correspondence between DTI microstructural indices and underlying myelin microstructure, where high FA was linked to high myelin density and a sharply tuned histological orientation profile, whereas high MD was related to diffuse histological orientation and low myelin density (Seehaus et al., 2015). Diffusion MRI studies applying more advanced biophysical models of white matter microstructure may be able to provide additional insight into the specific biological attributes underlying these brain-behaviour associations (Assaf et al., 2017; Huber et al., 2018).

The causes of inter-individual variation in white matter microstructure are not fully understood, but likely involve a complex interplay between genetic and environmental factors over the lifespan. Evidence from both adults and neonates, for instance, suggests that the microstructure of the fornix is highly heritable (Lee et al., 2015; Budisavljevic et al., 2016). The fornix is also one the earliest white matter tracts to mature, reaching its peak FA and minimum MD before age 20 (Lebel et al., 2012), and potentially nearing maturation during infancy and childhood (Dubois et al., 2008). At the same time, evidence suggests that fornix microstructure displays learning-related plasticity, even over short time periods. For instance, short-term spatial learning, in both rodents and humans, has been shown to induce alterations in diffusion indices of fornix microstructure (Hofstetter et al., 2013). Similarly, navigational ability is influenced by both genetic factors and experience (Lee and Spelke, 2010; Wolbers and Hegarty, 2010). Thus, fornix microstructure is likely to both shape, and be shaped by spatial navigation, in a bidirectional fashion (Bechler et al., 2018).

To conclude, by modelling learning performance on a virtual-reality water maze, we showed that the microstructure of the main white matter pathway linking the hippocampus and medial diencephalon – the fornix – predicted individual differences in human navigational learning. These results suggest that a full understanding of the biological underpinnings of individual differences in human navigational ability requires not only the analysis of individual processing regions, but of a distributed “navigation system”, underpinned by white matter. Critically, given the vulnerability of this brain system to the deleterious effects of aging (Lester et al., 2017), but also pathology in Alzheimer’s disease (Braak and Braak, 1991; Oishi et al., 2012), it is a key priority to develop behavioural markers of navigational ability that are sensitive to individual variation in this network, as seen here. One study in rodents, for instance, found that poorer learning on the MWM in early life predicted cognitive impairment in later life, but also that extensive training in poorer learners buffered against age-related learning impairments (Hullinger and Burger, 2015). Studies such as this highlight the potential of navigational learning, particularly as assessed using translation paradigms (Possin et al., 2016), for characterising, and potentially ameliorating, the effects of cognitive decline.

## Author contributions

CJH, MS, BK, ADE and KSG contributed to the conception and design of the experiment; MS collected imaging and behavioural data; CJH, MS, ANW, BK and JZ analysed the data; CJH wrote the manuscript with input from all other authors.

## Acknowledgments

We would like to thank Ofer Pasternak and Greg Parker for providing the free water correction pipeline, and John Evans and Peter Hobden for scanning support.

## References

Aggleton JP, O’Mara SM, Vann SD, Wright NF, Tsanov M, Erichsen JT (2010) Hippocampalanterior thalamic pathways for memory: Uncovering a network of direct and indirect actions. Eur J Neurosci 31:2292–2307.

Alonso JR, Hoi SU, Amaral DG (1996) Cholinergic innervation of the primate hippocampal formation: II. Effects of fimbria/fornix transection. J Comp Neurol 375:527–551.

Assaf Y, Johansen-Berg H, Thiebaut de Schotten M (2017) The role of diffusion MRI in neuroscience. NMR Biomed: 1–16.

Bechler ME, Swire M, ffrench-Constant C (2018) Intrinsic and adaptive myelination— A sequential mechanism for smart wiring in the brain. Dev Neurobiol 78:68–79.

Bohbot VD, Kalina M, Stepankova K, Spackova N, Petrides M, Nadel L (1998) Spatial memory deficits in patients with lesions to the right hippocampus and to the right parahippocampal cortex. Neuropsychologia 36:1217–1238.

Braak H, Braak E (1991) Neuropathological stageing of Alzheimer-related changes. Acta Neuropathol 82:239–259.

Buckley MJ, Wilson CRE, Gaffan D (2008) Fornix transection Iipairs visuospatial memory acquisition More than retrieval. Behav Neurosci 122:44–53.

Budisavljevic S, Kawadler JM, Dell’Acqua F, Rijsdijk F V., Kane F, Picchioni M, McGuire P, Toulopoulou T, Georgiades A, Kalidindi S, Kravariti E, Murray RM, Murphy DG, Craig MC, Catani M (2016) Heritability of the limbic networks. Soc Cogn Affect Neurosci 11:746–757.

Cain DP, Boon F, Corcoran ME (2006) Thalamic and hippocampal mechanisms in spatial navigation: A dissociation between brain mechanisms for learning how versus learning where to navigate. Behav Brain Res 170:241–256.

Chersi F, Burgess N (2015) The cognitive architecture of spatial navigation: hippocampal and striatal contributions. Neuron 88:64–77.

Chrastil ER, Sherrill KR, Aselcioglu I, Hasselmo ME, Stern CE (2017) Individual differences in human path integration abilities correlate with gray matter volume in retrosplenial cortex, hippocampus, and medial prefrontal cortex. Eneuro 4:ENEURO.0346-16.2017.

Concha L, Gross DW, Beaulieu C (2005) Diffusion tensor tractography of the limbic system. Am J Neuroradiol 26:2267–2274.

Croux C, Dehon C (2010) Influence functions of the Spearman and Kendall correlation measures. Stat Methods Appl 19:497–515.

De Bruin JPC, Moita MP, De Brabander HM, Joosten RNJMA (2001) Place and response learning of rats in a Morris water maze: Differential effects of fimbria fornix and medial prefrontal cortex lesions. Neurobiol Learn Mem 75:164–178.

Devan BD, Goad EH, Petri HL (1996) Dissociation of hippocampal and striatal contributions to spatial navigation in the water maze. Neurobiol Learn Mem 66:305–323.

Diedenhofen B, Musch J (2015) Cocor: A comprehensive solution for the statistical comparison of correlations. PLoS One 10:1–12.

Dubois J, Dehaene-Lambertz G, Perrin M, Mangin JF, Cointepas Y, Duchesnay E, Le Bihan D, Hertz-Pannier L (2008) Asynchrony of the early maturation of white matter bundles in healthy infants: Quantitative landmarks revealed noninvasively by diffusion tensor imaging. Hum Brain Mapp 29:14–27.

Dumont JR, Amin E, Wright NF, Dillingham CM, Aggleton JP (2015) The impact of fornix lesions in rats on spatial learning tasks sensitive to anterior thalamic and hippocampal damage. Behav Brain Res 278:360–374.

Eichenbaum H (2017) On the integration of space, time, and memory. Neuron 95:1007–1018.

Eichenbaum H, Stewart C, Morris RG (1990) Hippocampal representation in place learning. J Neurosci 10:3531–3542.

Ekstrom AD (2015) Why vision is important to how we navigate. Hippocampus 25:731–735.

Ekstrom AD, Huffman DJ, Starrett M (2017) Interacting networks of brain regions underlie human spatial navigation: A review and novel synthesis of the literature. J Neurophysiol:jn.00531.2017.

Ekstrom AD, Ranganath C (2017) Space, time, and episodic memory: The hippocampus is all over the cognitive map. Hippocampus:1–8.

Fidalgo C, Martin CB (2016) The hippocampus contribues to allocentric spatial memory through coherent scene representations. J Neurosci 36:2555–2557.

Gaffan D (1991) Spatial organization of episodic memory. Hippocampus 1:262–264.

Gaffan D (1992) Amnesia for complex naturalistic scenes and for objects following fornix transection in the rhesus monkey. Eur J Neurosci 4:381–388.

Gaffan D (1994) Scene-specific memory for objects: a model of episodic memory impairment in monkeys with fornix transection. J Cogn Neurosci 6:305–320.

Gaffan EA, Bannerman DM, Warburton EC, Aggleton JP (2001) Rats’ processing of visual scenes: Effects of lesions to fornix, anterior thalamus, mamillary nuclei or the retrohippocampal region. Behav Brain Res 121:103–117.

Hodgetts CJ, Postans M, Shine JP, Jones DK, Lawrence AD, Graham KS (2015) Dissociable roles of the inferior longitudinal fasciculus and fornix in face and place perception. Elife 4:e07902.

Hodgetts CJ, Postans M, Warne N, Varnava A, Lawrence AD, Graham KS (2017) Distinct contributions of the fornix and inferior longitudinal fasciculus to episodic and semantic autobiographical memory. Cortex 94:1–14.

Hofstetter S, Tavor I, Tzur Moryosef S, Assaf Y (2013) Short-Term Learning Induces White Matter Plasticity in the Fornix. J Neurosci 33:12844–12850.

Huber E, Henriques RN, Owen JP, Rokem A, Yeatman JD (2018) Applying biophysical models to understand the role of white matter in cognitive development. bioRxiv:347872.

Hullinger R, Burger C (2015) Learning impairments identified early in life are predictive of future impairments associated with aging. Behav Brain Res 294:224–233.

Iaria G, Barton JJS (2010) Developmental topographical disorientation: A newly discovered cognitive disorder. Exp Brain Res 206:189–196.

Iaria G, Bogod N, Fox CJ, Barton JJS (2009) Developmental topographical disorientation: Case one. Neuropsychologia 47:30–40.

Ikonen S, McMahan R, Gallagher M, Eichenbaum H, Tanila H (2002) Cholinergic system regulation of spatial representation by the hippocampus. Hippocampus 12:386–397.

Jankowski MM, Ronnqvist KC, Tsanov M, Vann SD, Wright NF, Erichsen JT, Aggleton JP, O’Mara SM (2013) The anterior thalamus provides a subcortical circuit supporting memory and spatial navigation. Front Syst Neurosci 7:45.

Jeurissen B, Leemans A, Jones DK, Tournier JD, Sijbers J (2011) Probabilistic fiber tracking using the residual bootstrap with constrained spherical deconvolution. Hum Brain Mapp 32:461–479.

Kahn AE, Mattar MG, Vettel JM, Wymbs NF, Grafton ST, Bassett DS (2017) Structural pathways supporting swift acquisition of new visuomotor skills. Cereb Cortex 27:173–184.

Kolarik BS, Shahlaie K, Hassan A, Borders AA, Kaufman KC, Gurkoff G, Yonelinas AP, Ekstrom AD (2016) Impairments in precision, rather than spatial strategy, characterize performance on the virtual Morris Water Maze: A case study. Neuropsychologia 80:90–101.

Lakens D (2016) Why you don’t need to adjust your alpha level for all tests you’ll do in your lifetime. 20% Stat Available at: http://daniellakens.blogspot.com/2016/02/why-you-dont-need-to-adjust-you-alpha.html.

Landau B, Lakusta L (2009) Spatial representation across species: geometry, language, and maps. Curr Opin Neurobiol 19:12–19.

Latini F (2015) New insights in the limbic modulation of visual inputs: The role of the inferior longitudinal fasciculus and the Li-Am bundle. Neurosurg Rev 38:179– 190.

Lebel C, Gee M, Camicioli R, Wieler M, Martin W, Beaulieu C (2012) Diffusion tensor imaging of white matter tract evolution over the lifespan. Neuroimage 60:340– 352.

Lee SA, Spelke ES (2010) Two systems of spatial representation underlying navigation. Exp Brain Res 206:179–188.

Lee SJ, Steiner RJ, Luo S, Neale MC, Styner M, Zhu H, Gilmore JH (2015) Quantitative tract-based white matter heritability in twin neonates. Neuroimage 111:123–135.

Leemans A, Jones DK (2009) The B-matrix must be rotated when correcting for subject motion in DTI data. Magn Reson Med 61:1336–1349.

Lester AW, Moffat SD, Wiener JM, Barnes CA, Wolbers T (2017) The aging navigational system. Neuron 95:1019–1035.

Maguire E a, Frackowiak RS, Frith CD (1997) Recalling routes around london: activation of the right hippocampus in taxi drivers. J Neurosci 17:7103–7110.

Metzler-Baddeley C, Jones DK, Belaroussi B, Aggleton JP, O’Sullivan MJ (2011) Frontotemporal Connections in Episodic Memory and Aging: A Diffusion MRI Tractography Study. J Neurosci 31:13236–13245.

Miller VM, Best PJ (1980) Spatial correlates of hippocampal unit activity are altered by lesions of the fornix and entorhinal cortex. Brain Res 194:311–323.

Moran NF, Lemieux L, Kitchen ND, Fish DR, Shorvon SD (2001) Extrahippocampal temporal lobe atrophy in temporal lobe epilepsy and mesial temporal sclerosis. Brain 124:167–175.

Morris R (1984) Developments of a water-maze procedure for studying spatial learning in the rat. J Neurosci Methods 11:47–60.

Murray EA, Wise SP, Graham (2016) The evolution of memory systems. Oxford, UK: Oxford University Press.

Murray EA, Wise SP, Graham KS (2017) Representational specializations of the hippocampus in phylogenetic perspective. Neurosci Lett.

O’Keefe J, Nadel L (1976) The hippocampus as a cognitive map. Oxford: Clarendon Press.

O’Keefe J, Nadel L, Keightley S, Kill D (1975) Fornix lesions selectively abolish place learning in the rat. Exp Neurol 48:152–166.

Oishi K, Mielke MM, Albert M, Lyketsos CG, Mori S (2012) The fornix sign: A potential sign for alzheimer’s disease based on diffusion tensor imaging. J Neuroimaging 22:365–374.

Olton DS, Walker JA, Gage FH (1978) Hippocampal connections and spatial discrimination. Brain Res 139:295–308.

Packard MG, Hirsh R, White NM (1989) Differential effects of fornix and caudate nucleus lesions on two radial maze tasks: evidence for multiple memory systems. J Neurosci 9:1465–1472.

Packard MG, McGaugh JL (1992) Double dissociation of fornix and caudate nucleus lesions on acquisition of two water maze tasks: further evidence for multiple memory systems. Behav Neurosci 106:439–446.

Pasternak O, Sochen N, Gur Y, Intrator N, Assaf Y (2009) Free water elimination and mapping from diffusion MRI. Magn Reson Med 62:717–730.

Patenaude B, Smith SM, Kennedy D, Jenkinson M (2012) NIH Public Access. Neuroimage 56:907–922.

Pereira T, Burwell RD (2015) Using the spatial learning index to evaluate performance on the water maze. Behav Neurosci 129:533–539.

Possin KL, Sanchez PE, Anderson-Bergman C, Fernandez R, Kerchner GA, Johnson ET, Davis A, Lo I, Bott NT, Kiely T, Fenesy MC, Miller BL, Kramer JH, Finkbeiner S (2016) Cross-species translation of the Morris maze for Alzheimer’s disease. J Clin Invest 126:779–783.

Postans M, Hodgetts CJ, Mundy ME, Jones DK, Lawrence AD, Graham KS (2014) Interindividual variation in fornix microstructure and macrostructure is related to visual discrimination accuracy for scenes but not faces. J Neurosci 34:12121– 12126.

Qi Z, Han M, Garel K, San Chen E, Gabrieli JDE (2015) White-matter structure in the right hemisphere predicts Mandarin Chinese learning success. J Neurolinguistics 33:14–28.

Ripollés P, Biel D, Peñaloza C, Kaufmann J, Marco-Pallarés J, Noesselt T, Rodríguez-Fornells A (2017) Strength of temporal white matter pathways predicts semantic learning. J Neurosci 37:1720–17.

Rudebeck SR, Scholz J, Millington R, Rohenkohl G, Johansen-Berg H, Lee ACH (2009) Fornix microstructure correlates with recollection but not familiarity memory. J Neurosci 29:14987–14992.

Ryan L, Lin CY, Ketcham K, Nadel L (2010) The role of medial temporal lobe in retrieving spatial and nonspatial relations from episodic and semantic memory. Hippocampus 20:11–18.

Saunders RC, Aggleton JP (2007) Origin and topography of fibers contributing to the fornix in macaque monkeys. Hippocampus 17:396–411.

Schinazi VR, Nardi D, Newcombe NS, Shipley TF, Epstein RA (2013) Hippocampal size predicts rapid learning of a cognitive map in humans. Hippocampus 23:515–528.

Shapiro ML, Simon DK, Olton DS, Gage FH, Nilsson O, Björklund A (1989) Intrahippocampal grafts of fetal basal forebrain tissue alter place fields in the hippocampus of rats with fimbria-fornix lesions. Neuroscience 32:1–18.

Simpson EL, Gaffan EA, Eacott MJ (1998) Rats’ object-in-place encoding and the effect of fornix transection. Psychobiol 26:190–204.

Steiger JH (1980) Tests for comparing elements of a correlation matrix. Psychol Bull 87:245–251.

Stepanov II, Abramson CI (2008) The application of the first order system transfer function for fitting the 3-arm radial maze learning curve. J Math Psychol 52:311– 321.

Sutherland RJ, Rodriguez AJ (1989) The role of the fornix/fimbria and some related subcortical structures in place learning and memory. Behav Brain Res 32:265– 277.

Tolman EC (1948) Cognitive maps in rats and men. Psychol Rev 55:189–208.

Tuch DS, Reese TG, Wiegell MR, Makris N, Belliveau JW, Van Wedeen J (2002) High angular resolution diffusion imaging reveals intravoxel white matter fiber heterogeneity. Magn Reson Med 48:577–582.

Wakana S, Caprihan A, Panzenboeck MM, Fallon JH, Perry M, Gollub RL, Hua K, Zhang J, Jiang H, Dubey P, Blitz A, van Zijl P, Mori S (2007) Reproducibility of quantitative tractography methods applied to cerebral white matter. Neuroimage 36:630–644.

Warburton EC, Aggleton JP (1998) Differential deficits in the Morris water maze following cytotoxic lesions of the anterior thalamus and fornix transection. Behav Brain Res 98:27–38.

Warburton EC, Aggleton JP, Muir JL (1998) Comparing the effects of selective cingulate cortex lesions and cingulum bundle lesions on water maze performance by rats. Eur J Neurosci 10:622–634.

Westman E, Aguilar C, Muehlboeck JS, Simmons A (2013) Regional magnetic resonance imaging measures for multivariate analysis in Alzheimer’s disease and mild cognitive impairment. Brain Topogr 26:9–23.

Wetzels R, Wagenmakers EJ (2012) A default Bayesian hypothesis test for correlations and partial correlations. Psychon Bull Rev 19:1057–1064.

Whishaw IQ, Maaswinkel H (1998) Rats with fimbria-fornix lesions are impaired in path integration: a role for the hippocampus in “sense of direction”. J Neurosci 18:3050–3058.

Wilcox RR (2016) Comparing dependent robust correlations. Br J Math Stat Psychol 69:215–224.

Wolbers T, Hegarty M (2010) What determines our navigational abilities? Trends Cogn Sci 14:138–146.

Wolbers T, Wiener JM (2014) Challenges for identifying the neural mechanisms that support spatial navigation: the impact of spatial scale. Front Hum Neurosci 8:571.

